# The interplay of active and passive mechanisms in slow axonal transport

**DOI:** 10.1101/2022.01.23.477383

**Authors:** Reshma Maiya, Swagata Dey, Krishanu Ray, Gautam I. Menon

## Abstract

A combination of intermittent active movement of transient aggregates and a paused state that intervenes between periods of active transport has been proposed to underly the slow, directed transport of soluble proteins in axons. A component of passive diffusion in the axoplasm may also contribute to slow axonal transport, although quantitative estimates of the relative contributions of diffusive and active movement in the slow transport of a soluble protein, and in particular how they might vary across developmental stages, are lacking. Here, we propose and study a model for slow axonal transport, addressing data from bleach-recovery measurements on a small, soluble, protein, Choline Acetyltransferase (ChAT), in thin axons of the *lateral chordotonal* (*lch5*) sensory neurons of *Drosophila*. ChAT is mainly present in soluble form in the axon and catalyses the acetylation of choline at the synapse. It does not form particulate structures in axons and moves at rates characteristic of slow component b (≈ 1-10 mm/day or 0.01-0.1 *μm/s*). Using our model, which incorporates active transport, paused and diffusive states, we predict bleach recovery and cargo trajectories obtained through kymographs, comparing these to experimental observations at different developmental stages. We show that changes in the diffusive fraction of cargo during these developmental stages dominate bleach recovery and that a combination of active motion with a paused state alone cannot reproduce the data. We compared predictions of the model with results from photoactivation experiments. The importance of the diffusive state in reproducing the bleach recovery signal in the slow axonal transport of small soluble proteins is our central result.

**STATEMENT OF SIGNIFICANCE:** While the fast axonal transport of cargo in axons is by now well-understood, the nature of slow transport remains controversial. A number of different models having been proposed for slow axonal transport, including models which allow for transitions between an intermittently moving molecular-motor driven state and a stalled state. How mechanisms for slow axonal transport are modulated during development is unexplored. We study a number of different models for slow axonal transport, comparing their predictions to data on transport of the enzyme Choline Acetyltransferase (ChAT) in thin *lateral chordotonal* (*lch5*) sensory neurons of *Drosophila* larva, across developmental stages where flux increases significantly. We show that accounting for changes in the diffusive fraction of cargo during these developmental stages is essential and diffusion cannot be neglected in the modelling of the slow axonal transport of small soluble proteins.

## INTRODUCTION

Axonal transport of cellular cargo is largely driven through the activity of molecular motors. These motors move on the highly polarized microtubule tracks that crowd the axon. Cargo moving away from the cell body and towards the synapse is said to move in the anterograde direction, whereas cargo returning towards the cell body moves retrogradely. Kinesin-1 (1, 2) and Kinesin-2 (3) are among the motors driving anterograde transport, while Dynein drives retrograde transport (2). Motor-driven cargo typically exhibit bidirectional movement, often pausing or reversing their direction of motion. This indicates that different motor proteins can bind the same cargo simultaneously, with tug-of-war mechanisms (4–6) or selective regulation of specific motor types (7) determining transport properties.

Axonal transport is categorised into two rate components, fast and slow (8, 9). Membrane-bound organelles and vesicles move at velocities of 0.6 – 5 *μm s*^−1^, defining the fast component. Cytoskeletal proteins in oligomeric form, as well as some small soluble proteins, are transported at slower velocities of 0.002 – 0.09 *μm s*^−1^. This defines the slow component of axonal transport (10–12). The slow component further subsumes two rate components (a and b), termed SCa and SCb respectively, depending on the nature of the cargo being transported. Cytoskeletal elements, including tubulin, neurofilaments, spectrin and tau proteins, typically constitute the SCa rate component. Actin, Clathrin and Calmodulin, as well as metabolic enzymes such as Glyceraldehyde-3-phosphate dehydrogenase (GAPDH), inactive Dynein, Choline acetyltransferase (ChAT), Synapsin and *α*-Synuclein (8, 13), are transported through the SCb route. SCa rate components move at about 0.2 – 1 mm/day whereas SCb components move at speeds of about 2 – 8 mm/day (10).

Though SCa and SCb components can be seen to move intermittently at fast component rates, their overall motion is dominated by diffusion or pauses giving rise to slower transport rates on average. Prominent models for slow transport are the “stop and go” model, the “dynamic recruitment” model and the “Kinesin limited” model (11, 14–16). In the “stop and go” model for the slow transport of neurofilaments, movement occurs through short, intermittent bursts of directed motion interspersed with long pauses (14, 17). Transport is slow overall, because neurofilament cargo spends significant amount of time in the paused state as well as in moving in the opposite direction. Other models describe the transport of specific cytosolic proteins. The “dynamic recruitment” model (15), is based on observations of Synapsin and CAMIIK, both of which exhibit affinity towards membranes. The “dynamic recruitment” model proposes that these proteins can transiently assemble into larger, dynamic complexes. These can then be transported by motors when they attach to a moving membrane-bound vesicle. The recruitment of these complexes results in their intermittent active transport. This process is interrupted by the disassembly of the aggregate and/or the detachment of the motor. Finally, the “Kinesin-limited” model (16) explains the vesicle-independent anterograde transport of inactive Dynein by Kinesin-1. This association is less stable and limited by competitive binding of activated Kinesin-1 to other cargo, accounting for the name.

All these models make some basic assumptions: The first, common to all, is that motion of the slow component is sporadic. There are long periods where the cargo is not bound to an active motor involved in translocation. The second, assumed in some models but not in others, is that cargo transported by this route may be aggregated, though aggregates need not be motor bound. Such aggregates appear as particulate structures in imaging experiments. The third assumption deals with the behaviour in-between motor-driven transport events: most models implicitly assume that cargo are stationary or paused on-track. Fourth, the assumption is that the amount of motor present should be a limiting factor in slow transport, since competitive binding to cargo transported by a fast route reduces the amount of motor proteins available to take up the slow component. To what extent the situations described by these different models might overlap is unclear.

How these mechanisms might be differentially modulated across developmental stages is unclear. Measurements of the transport of Choline Acetyltransferase (ChAT), a small soluble enzyme synthesised in the cell body and found enriched at presynaptic terminals in *Drosophila*, can potentially shed light on these questions. ChAT is found in both membrane bound and soluble form. Around 80% of ChAT in Drosophila is found in soluble form while the rest is membrane bound. In membrane-bound form, it is associated with synaptic vesicles (18, 19). Sadananda et al.(13) have examined the transport of ChAT in the axons of *Drosophila*, observing ChAT moving in association with Kinesin-2, but without the formation of particulate structures. They show that during specific developmental stages in the third instar larva, ChAT molecules move at relatively faster overall rates. This transport is driven by the heterotrimeric Kinesin-2 motor.

Dey and Ray (20) used Fluorescence Recovery after Photobleaching (FRAP) methods to study the transport characteristics of ChAT and Kinesin-2 motor subunits over four developmental stages during 76h - 79h After Egg Laying (AEL) in *lateral chordotonal* (*lch5*) neurons. FRAP is a live-cell imaging technique that involves expressing the molecules of interest with fluroscent tags which are photobleached irresversibly in a small region and the movement of unbleached fluorescent molecules from the neighbourhood into the bleached region is followed (21). ChAT intensity increases four-fold while the intensity of Kinesin-2 motor increases by less than a factor of 2. The paradox posed by the data is the following: Across developmental stages 76-78h AEL, one sees an increase in cargo, in motors and in cargo-motor interactions. Sensitized Forster’s Resonance Energy Transfer (sFRET) assay was used to study the interaction between ChAT and Kinesin-2 motor subunits.The technique is used to study the dynamics of the protein or cell components of interest, their localization and interactions with other cellular components (22). sFRET measurements between TQ-ChAT and KLP68D-YFP indicate a 10-20% increase of interaction between cargo and motor from 76h to 78h AEL (Figure 1 A). An increased number of motors and of cargo-motor interaction would favour an active component to transport, but that must be set off against an increase in the total amount of cargo.

**Figure 1:**
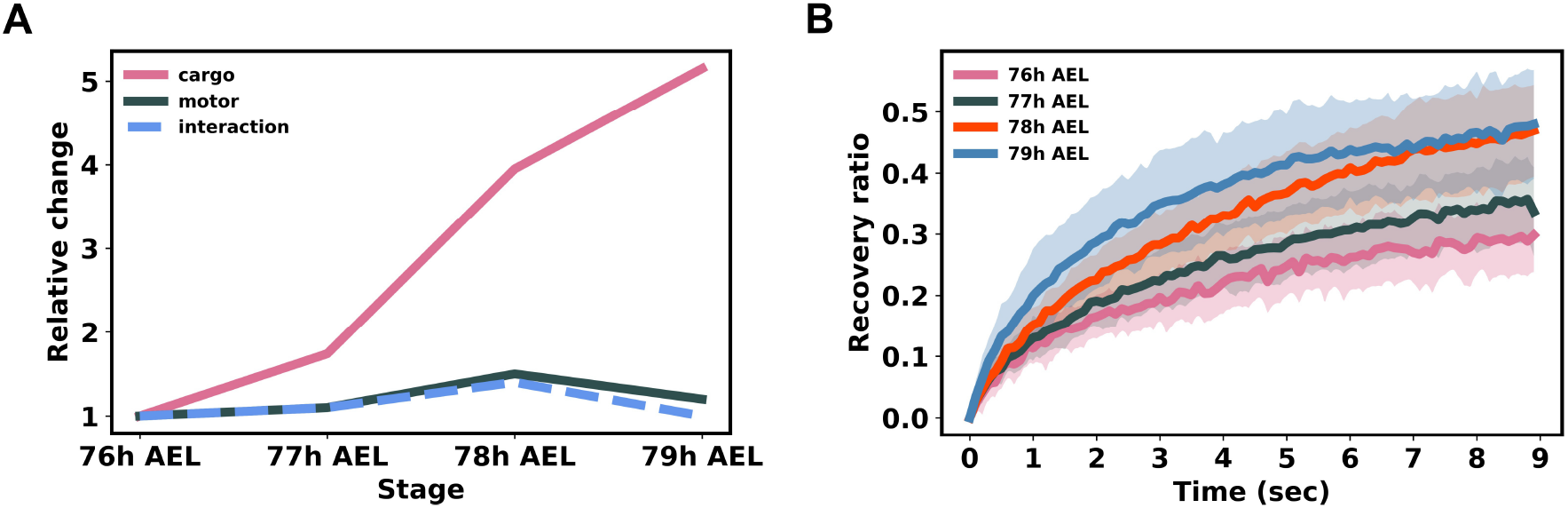
Experimental data on (A) changes in the levels of cargo, motor and interaction between cargo and motor relative to those at 76h AEL, (B) bleach recovery profiles across developmental stages 76-79h AEL (Data taken from (20))

Here, we propose and study several models for bleach recovery of ChAT across the 76h - 79h AEL in *Drosophila*, testing our model predictions against data from Dey and Ray (20). We perform kinetic Monte-Carlo simulations of these models, using them to describe the interplay of active and diffusive transport of ChAT. We show that accounting for diffusion is crucial to interpreting FRAP data on slow axonal transport and that the models that account for only the paused and active transport states cannot reproduce the results of the bleach recovery experiments across developmental stages. Accounting for diffusion that intervenes in-between periods of active transport provides an accurate biophysical description of this problem, consistent with data for cargo and motor levels, as well as motor-cargo interactions. We found that the increase in cargo *outweighed* the increase in both motor numbers and in motor-cargo interactions, allowing a larger fraction of cargo to be transported by passive diffusion. The comparisons we make to data provide a detailed picture of how the interaction of diffusive and active transport might be tuned across developmental stages. They also provide methods applicable in similar contexts across other model systems.

## METHODS

We idealize the bundle of highly-polarized microtubules in the axon as a single effective filament on which cargo is actively transported by molecular motors. This averages over the circumferential dimension of the axonal microtubule bundle, a reasonable approximation given the narrowness of the axon, but which requires the use of an effective one-dimensional diffusion constant. This filament is represented by a one-dimensional lattice (Figure 2 B). Cargo moves along this filament when it is attached to a motor. ChAT proteins are modelled as particles moving at fixed velocity on a filament when bound to a motor. When cargo is disengaged from the track, we assume that it can freely diffuse within the confines of the axoplasm (Figure 2 C and D) or remain paused. Interactions between cargo particles are not considered. We tune rates for both active and the diffusive cargo so as to reproduce numbers, such as diffusion constants and hopping rates, that are extracted from the experimental data, inferred from biophysical arguments or derived from previous work.

**Figure 2:**
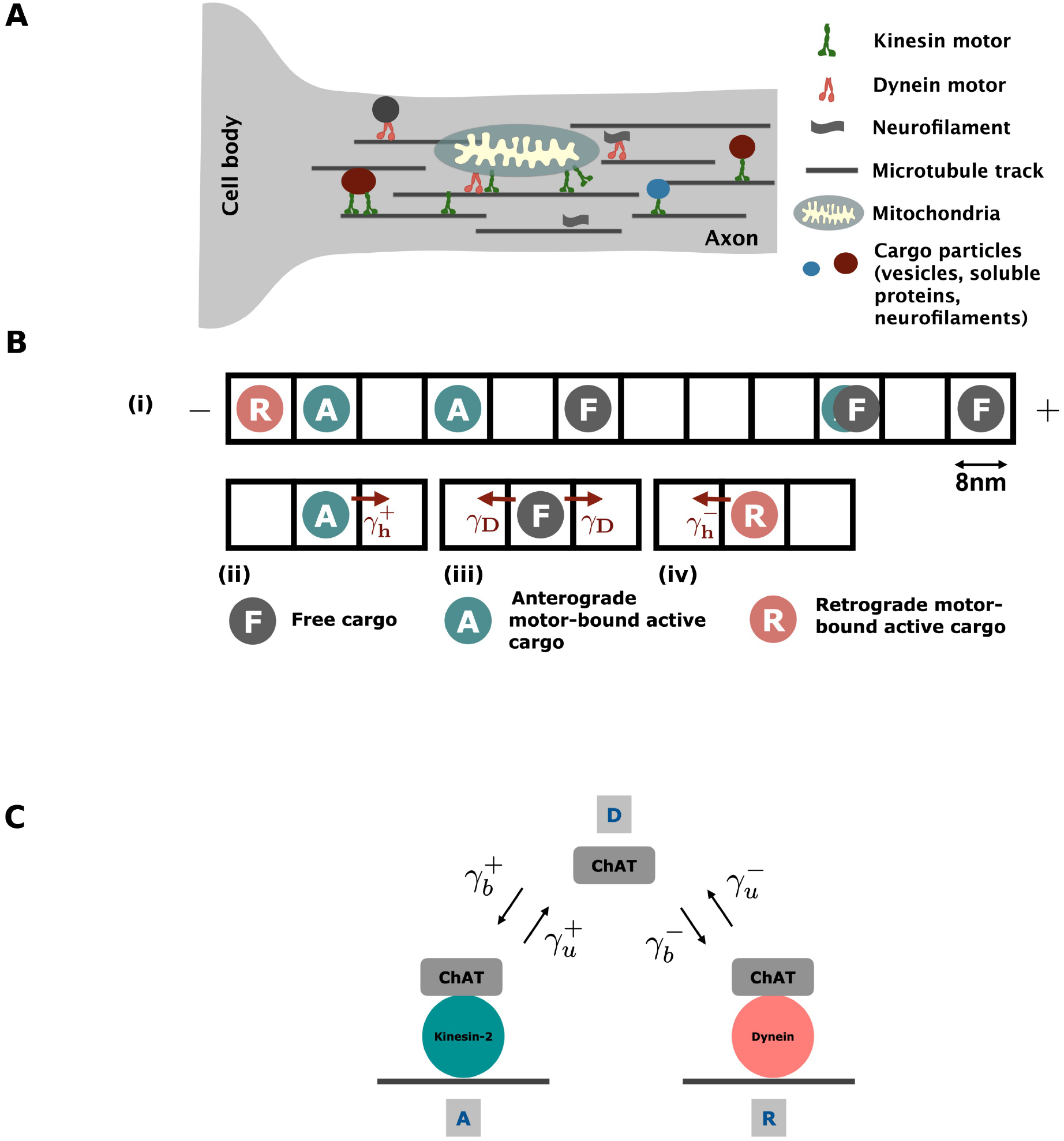
Schematic of (A) the axon showing the variety of cargo being transported, the bundle of axonal microtubules and a large cargo, a single mitochondrion. (B) (i) Lattice representing the bundle of microtubules with the lattice sites spaces 8nm apart. Occupancy of each site on the lattice is the sum total of cargo occupancies on all microtubule tracks at that distance along the axon. Diffusion and active hopping motions of cargo in diffusive (iii) and active (ii,iv) states. (C) The three state model for motor-driven axonal transport, depicting 3 states for the motor-cargo complex - two active states and one diffusive or paused state, transitions between these states with the associated rates. These are described in the text.

We describe a number of models of differing complexity. The first is a model that accounts for motor-driven - anterograde and retrograde - transport, in addition to a paused state (Figure 2 C). This is most analogous to the ‘stop and go’ model, initially introduced for neurofilament transport. We then go on to discuss a more general 3-state model, which includes a diffusive state in addition to motor-driven active transport states. Finally, we describe a model that has actively moving, paused and diffusive states, a 5-state model (Figure S1); results for this model are relegated to the Supplementary Information.

Specifying these models require specifying a large number of rates and relative fractions of diffusive and active components. The rates to be specified include hopping rates, both anterograde and retrograde, motor binding and unbinding rates and a number of diffusion constants. We benchmark these with the limited experimental data available, through biophysical estimates as well as fits to the data itself.

### Model definitions

#### Three state model with a paused state

The three state model with a paused state accounts for the following cargo motion states: (I) an anterograde motor-bound active state, (II) a retrograde motor-bound active state and (III) a paused state. As shown in the schematic of Figure 2 C, in an infinitesimal time interval a particle can undergo one of the several of the following processes: A particle in the anterograde state can move one site to the right, at a rate 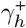 (Figure 2 B (ii)), while a particle in the retrograde state can move one site to the left, at a rate 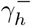 (Figure 2 B (iv)) reproducing the velocities of anterograde and retrograde motors. A particle in either of these states can switch to becoming a paused particle at rates 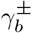 (Figure 2 C). The paused state is described by setting the diffusion constant in that state, *γ_D_* = 0. A paused particle can reconvert to either a retrogradely or anterogradely moving particle at rates 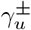 (Figure 2 C). The fundamental rates used in our simulation are listed in Table 1. Wherever possible we use rates extracted from literature and our own estimates for those rates for which prior information is unavailable. These rates reflect underlying processes of motor-cargo attachment, the productive encounters of motor-cargo complexes with the filament and the transition between bound and unbound states.

**Table 1:** Parameters common to all models.

| Symbol | Parameter | Value |
| --- | --- | --- |
| $v_+$ | velocity of Kinesin-2 motor (29, 30) | $0.3\mu ms^{-1}$ |
| $v_-$ | velocity of Dynein motor (24) | $0.68\mu ms^{-1}$ |
| $l_+$ | mean run-length of Kinesin-2 motor (30) | $1.14\mu m$ |
| $l_-$ | mean run-length of Dynein motor (24) | $0.90\mu m$ |
| $h$ | step-length on the microtubule track | $8nm$ |
| $T$ | temperature | $296K$ |
| $\eta$ | viscosity of cytoplasm (13) | $3 \times 10^{-3} Pa.s$ |
| $r^{GFP::ChAT}$ | hydrodynamic radius of GFP::ChAT (13) | $5.79nm$ |
| $D$ | bulk diffusivity of cargo <sup>a</sup> | $1.2\mu m^2 s^{-1}$ |
| $a$ | radius of axon | $0.25\mu m$ |
| $\gamma_h^+$ | hopping rate of Kinesin-2 bound anterograde particle <sup>b</sup> | $0.0375 \times 10^3 s^{-1}$ |
| $\gamma_h^-$ | hopping rate of Dynein bound retrograde particle <sup>b</sup> | $0.085 \times 10^3 s^{-1}$ |
| $\tilde{\gamma}_u^+$ | unbinding rate of Kinesin-2 bound anterograde particle <sup>c</sup> | $0.2603 s^{-1}$ |
| $\tilde{\gamma}_u^-$ | unbinding rate of Dynein bound retrograde particle <sup>c</sup> | $0.755 s^{-1}$ |

<sup>a</sup> Estimated to be between 1-3 $\mu m^2/s$ <sup>b</sup> $v_{\pm}/h$ <sup>c</sup> $v_{\pm}/l_{\pm}$ Table 2: **Model specific parameters** (Estimated. Details in ‘Methods’)

#### Three state model with a diffusive state

The three state model with a diffusive state consists of (I) an anterograde motor-bound active state, (II) a retrograde motor-bound active state, (III) a diffusive state. Transitions between cargo states are as for the three state with the paused state involving transitions from the anterograde and retrograde active states to the diffusive state at rates 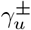 and from diffusive to anterograde and retrograde active states at rates 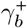 and 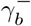 respectively (Figure 2 C). Cargo in the diffusive state moves with equal rate *γ_D_* in either direction.

A more detailed 5-state model is described in the Supplementary Information. Briefly, it admits the possibility of both motor-bound but paused states in addition to the diffusive states, interpolating between the models described above.

### Numerical methodology

#### Continuous-time Monte Carlo simulations

Once rates are specified, given an initial configuration of N particles, we update their positions using a continuous time Monte Carlo algorithm, the Gillespie algorithm (23). This algorithm splits each update into two steps. Given rates for the set of all possible distinct updates, say *r*_1_, *r*_2_ …, one first calculates the sum of these rates: 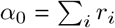. This is also called the total propensity. The time until the next update is then calculated as *τ* = (1/*α*_0_)(1/ϒ) where ϒ is a random number drawn from a uniform distribution in [0, 1]. In the second step, which specific update will occur in that time is computed by fully splitting the interval [0, 1] in proportion to the weighted ratio of rates. If the rate is larger, that corresponds to a larger subset of the range [0, 1]. A second random number is then called and, depending on its location in that interval, the system configuration is then updated accordingly and the procedure repeated. The time at which each update occurs is recorded. Each Gillespie step leads to an update of a single particle. Once *N* particles are updated, a single Monte Carlo step is said to have occurred.

#### Lattice model simulations

The system of cargo and microtubules is simulated on a lattice of size 3900 with successive lattice sites being 8 nm apart. This represents an axonal segment of length 31.2*μm*. The 8 nm lattice size corresponds to the modal step size of the motor proteins, given the distance between two successive *α – β* tubulin dimers on the microtubule lattice. We begin by randomly placing cargo particles in one of the cargo states across the lattice. Cargo states are chosen based on the estimated relative probabilities of finding them (see ‘Parameter estimation’). We checked with initially random assignments of cargo states that the same final proportions were obtained. We allow the system to settle into steady state by evolving this configuration across 10 seconds (10^7^ updates). The benchmarking of our rates in terms of physical time is provided in Table 1, 2.

**Table 2:** Model specific parameters. (Estimated. Details in ‘Methods’)

|  | Parameter | 76h<br>AEL | 77h<br>AEL | 78h<br>AEL | 79h<br>AEL |
| --- | --- | --- | --- | --- | --- |
| **3 state model without diffusion** |  |  |  |  |  |
| $\gamma_b^+(s^{-1})$ | rate of Kinesin-2-ChAT complex binding to microtubule track | 1.56 | 0.23 | 0.08 | 0.05 |
| $\gamma_b^-(s^{-1})$ | rate of Dynein-ChAT complex binding to microtubule track | 2.27 | 0.32 | 0.09 | 0.06 |
| **3 state model** |  |  |  |  |  |
| $\gamma_b^+(s^{-1})$ | rate of Kinesin-2-ChAT complex binding to microtubule track | 0.23 | 0.1 | 0.04 | 0.02 |
| $\gamma_b^-(s^{-1})$ | rate of Dynein-ChAT complex binding to microtubule track | 0.47 | 0.17 | 0.07 | 0.04 |

#### Bleaching protocol

Photobleaching simulations are carried out in a system with 100 particles. Since the bleach recoveries do not depend on the total cargo in the system but on the relative distribution of cargo in different cargo states, simulations for all developmental stages was carried out with 100 particles with different cargo distributions. To implement the bleaching protocol, the particles in the central 1/3rd of the simulated region (10.4*μm*) are tagged as “bleached”. This procedure mimics the photobleaching of particles in a FRAP experiment. The system is then allowed to evolve for another 9 seconds to permit comparisons with the transport behaviour for the duration in the FRAP experiments. The location, bleach status and the type of each cargo particle is tracked. Individual particle trajectories are recorded every 0.1 second corresponding to an experimental frame rate of 10 FPS.

We largely simulate periodic boundary conditions, in which the location of a particle at the last site moving one site to the right is updated to the first site and that of a particle in the first site moving a site to the left is updated to the last site. The methodology can be extended to open boundary conditions, in which case the input and output rates of the particles at the two terminal sites must also be specified, in addition to hopping rates in the bulk. Provided the system is large enough that bleached vesicles do not reenter the bleached region during the course of the simulation, the results are independent of the boundary condition and only depend on the average, steady state current which we have benchmarked.

From these data, we can extract bleach recovery profiles. Since we have access, in our simulations, to all particle coordinates, we can generate kymographs for unbleached particles. These are plotted by connecting the trajectories of individual unbleached particles sampled at the specified frame rate.

### Measurements and averaging

We compute the recovery ratio in the bleached region by taking the ratio of number of particles tagged to be bleached to the numbers before bleaching in the region. These recovery ratios are averaged over values from 100 simulations, each with a different starting system configuration and after the initial averaging described earlier. FRAP profile densities are plotted by taking the particle density at a site to be the sum of the number of particles over 100 simulations at that site and given time. The discrete simulation kymographs provide trajectories of individual particles. The qualitative features of these can be compared to those from the experimental kymographs.

### Parameter estimation

#### Hopping and unbinding rates

The hopping 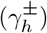 and the unbinding rates 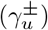 for cargo in active anterograde and retrograde states are calculated from experimental values of the average velocities (*v*_+_, *v*_−_) and mean run-lengths (*l*_+_, *l*_−_) of Kinesin-2 and Dynein motors respectively (24). Using the step-lengths of motors on microtubule tracks, defined as *h*, we then have 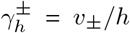 and 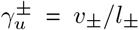. We also assume, as a reasonable first approximation that *unbinding* rates are independent of developmental stage.

#### Binding rates

Binding rates are computed using the equations describing the system (25) in steady state with periodic boundary conditions. For the 3-state model, solving the equations describing the system in steady state with periodic boundary conditions yields 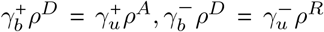 and 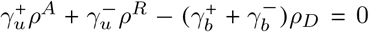. Normalizing to the total cargo *ρ*^T^ gives, 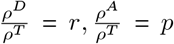 and 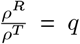. We therefore have binding rates in terms of the cargo fractions 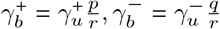.

To arrive at the relative distribution of cargo in different motion states, a certain amount of total (*ρ^T^* = *ρ^A^* + *ρ^R^* + *ρ^D^*) and active anterograde (*ρ^A^*) cargo is assumed at 76h AEL. Different combinations for values of amount of active anterograde (*ρ^A^*) and retrograde (*ρ^R^*) cargo at 76h AEL are tried to arrive at relative distributions of cargo in the three motion states 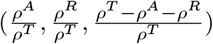 that give better fits to the experimental recovery curve. For cargo distributions in developmental stages 77-79h AEL, amounts of total (*ρ^T^*) and active anterograde (*ρ^A^*) cargo at 76h AEL are multiplied by a factor equal to change in the fluorescence intensity levels of TQ-ChAT and KLP68D at that stage relative to 76h AEL. In doing so, KLP68D levels are taken to be indicative of the amount of cargo bound to Kinesin-2 motors. The amount of active retrograde cargo in the system is assumed to be unchanging over the developmental stages (details in Table S3).

#### Diffusion rates

The bulk diffusivity (D) is calculated using the standard Stokes-Einstein expression, which relates the diffusion constant as proportional to the viscosity (*η*), hydrodynamic radius (*r*) of ChAT and absolute temperature(T) as 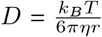, where *k_B_* is the Boltzmann constant with value 1.38 × 10^−23^ *JK*^−1^. Using standard values, the bulk diffusivity of ChAT turns out to be 12.467 *μm*^2^*s*^−1^ i.e. diffusion rate of 3.9 × 10^5^ *s*^−1^.

This is likely an overestimation for the following reasons. The conversion of a confined 3-d problem into an effective 1-d problem reduces the effective 1-d diffusion constant. Also the viscosity of the crowded axoplasm is larger (typically by a factor of 10 or more) than the viscosity of the water assumed above. We may conservatively take the diffusion constant to be smaller than the above number and use the reduced value of diffusion rate = 37500 *s*^−1^ (or diffusivity D = 1.2 *μm*^2^*s*^−1^) in our calculation for the diffusing states. Moreover, proteins of size 70 - 100 kDa in *E. coli* were estimated to have *D* ~ 1 – 2 *μm*^2^*s*^−1^ (26), while tau protein isoforms of size ranging from 68-100 kDa have diffusion constants of 3 *μm*^2^*s*^−1^ in neurons (27). Thus the diffusion constant for the 67 kDa ChAT molecule (28) can be expected to be in the range 1 – 3 *μm*^2^*s*^−1^.

Diffusion constant of the cargo in the paused states is taken to be zero in all the models.

### Photoactivation

#### Experimental photoactivation measurements

As an independent test of our model, we performed photoactivation experiments. The FRAP data from our previous study (20) indicated that Kinesin-2 motor is required for entry and accumulation of the bulk of ChAT in the axon and that it facilitates the anterograde progression of the ChAT inside the axon. The episodic movement of ChAT was further tested using a photoactivation assay, which involves selective photoactivation of fluorescence tag molecule attached to the protein or cell component of interest in a small region and tracking its motion (31). A variant of GFP (photoactivatable GFP, paGFP) that shows an increased fluorescence after being stimulated by 405nm wavelength was used for the photoactivation assay.

The photoactivation assay was carried out by expressing a GPAC tagged ChAT in lch5 neurons using chaGal4 and samples of age 76h-79h AEL were imaged. GPAC is a fusion fluorescent protein with photoactivatable GFP (paGFP) on the N-terminus and mCherry on the C-terminus (a kind gift from A. Welman, Edinburgh Cancer Research Centre). Axons spanning an area of 40.6 × 11.5 *μm*^2^ were imaged and photoactivation was carried out in an area of 8.05 *μm* (70 px) x 5.75 *μm* (50 px) in the cell body adjacent to the axon. The neurons were photoactivated by 0.52mW of 405nm laser and imaged by 488nm and 561nm for GFP and mCherry acquisition. Subsequently, movements of activated GPAC-ChAT fluorescence into the axons were tracked.

Immediately after activation in the cell bodies, the flow of activated GFP-ChAT was observed in the axons. The intensity of GFP-ChAT was quantified before and after photoactivation in a 5 *μm* region (proximal) of the axon. Axonal inflow was measured as increase in the axonal fluorescence (I_post-activation_ – I_pre-activation_) normalized to the intensity before photoactivation (I_pre-activation_).

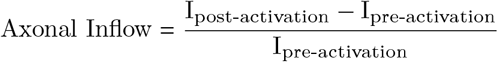

#### Photoactivation simulation

Photo-activation is simulated on a 35*μm* region with 50,87 and 197 cargo particles for 76, 77 and 78h AEL (following the trend in TQ-ChAT levels across developmental stages 76 to 79h AEL relative to at 76h AEL). The central 5*μm* region is taken to be the proximal AIS where the measurements are carried out. Once the system settles, cargo in the proximal 15*μm* region are marked to be ‘photo-activated’ and the simulation is run for 1s. The amount of cargo labelled ’photo-activated’ in the 5*μm* proximal AIS region is noted down after 1s.

## RESULTS & DISCUSSION

### Bleach recovery profiles across developmental stages: 3-state model with a paused state

To test if diffusion is crucial in the transport of ChAT, we first simulate the system in the absence of diffusion. Our results for the 3 state model with a paused state are shown in Figure 3. In Figure 3(A) we display experimentally derived colour kymographs from (20) showing the evolution of ChAT densities in the bleached region of the axon. In Figure 3(B) we show simulation predictions for this recovery. Particle kymographs across a larger simulation region encompassing the bleached region are shown in Figure 3(C).

**Figure 3:**
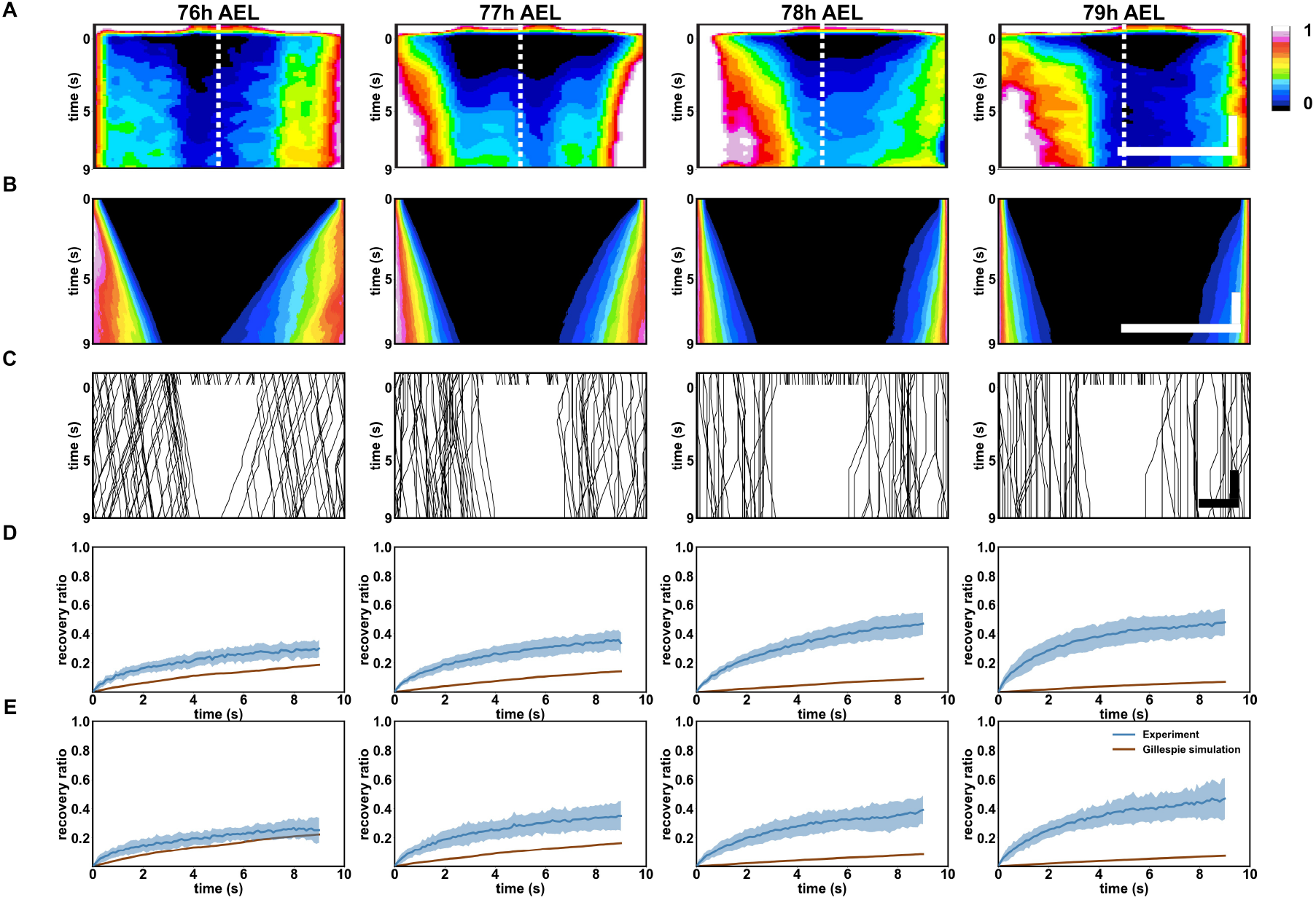
Results for the three state model with a paused state:(A-B)Colour kymographs of total cargo densities in the bleached region over 9 seconds from (A) experiment [(20) Image reproduced with permission from ..] and (B) Gillespie simulations for stages 76 - 79h AEL, (C) line kymographs over the entire region simulated using Gillespie algorithm for one run of the simulation, (D) recovery profiles in the bleached region from experiments and Gillespie simulations compared over 9 seconds. Scale bar - x-axis: 5*μm*, y-axis: 2*s*

The total fluorescent recovery in the bleached region over time is plotted as the fraction of the pre-bleach fluorescence in the bleached region in Figure 3(D) across the developmental stages 76h - 79h AEL with error bars. It can be seen that the bleach recoveries are much slower than in the experiments (Figure 3 B-D). The bleach recovery curves are also lower in successive developmental stages owing to decrease in the fraction of active cargo that is the sole contributor to bleach recovery (Table 3). This shows that active transport with interspersed stationary states cannot produce the bleach recovery profiles seen in experiments. The bleach recovery curves (Figure 3(D)) also rise linearly with time without exhibiting the gradual rise and tendency to saturate that is seen in the experiments.

**Table 3:** Cargo distribution (%) in 3 states for the parameter used in the Gillespie simulation.

| Developmental Stage | Active anterograde (p) | Active retrograde (q) | Paused/Diffusive (r) |
| --- | --- | --- | --- |
| **3 state model with a paused state** |  |  |  |
| 76h | 60 | 30 | 10 |
| 77h | 39 | 18 | 43 |
| 78h | 22 | 8 | 70 |
| 79h | 14 | 6 | 80 |
| **3 state model with a diffusive state** |  |  |  |
| 76h | 35 | 25 | 40 |
| 77h | 23 | 14 | 63 |
| 78h | 13 | 7 | 80 |
| 79h | 8 | 5 | 87 |

### Bleach recovery profiles across developmental stages: 3-state model with a diffusive state

Results from our simulations of a 3 state model in which one of the states is a diffusive state are shown in Figure 4. Experimentally derived colour kymographs from (20) (Figure 4(A)) depicting the evolution of ChAT densities in the bleached region of the axon is compared with those from the Gillespie simulation (Figure 4(B)). we show predictions for this recovery. Particle kymographs across a larger simulation region encompassing the bleached region are shown in Figure 4(C). The total fluorescent recovery in the bleached region over time is plotted as the fraction of the pre-bleach fluorescence in the bleached region in Figure 4(D) across the developmental stages 76h - 79h AEL with error bars.

**Figure 4:**
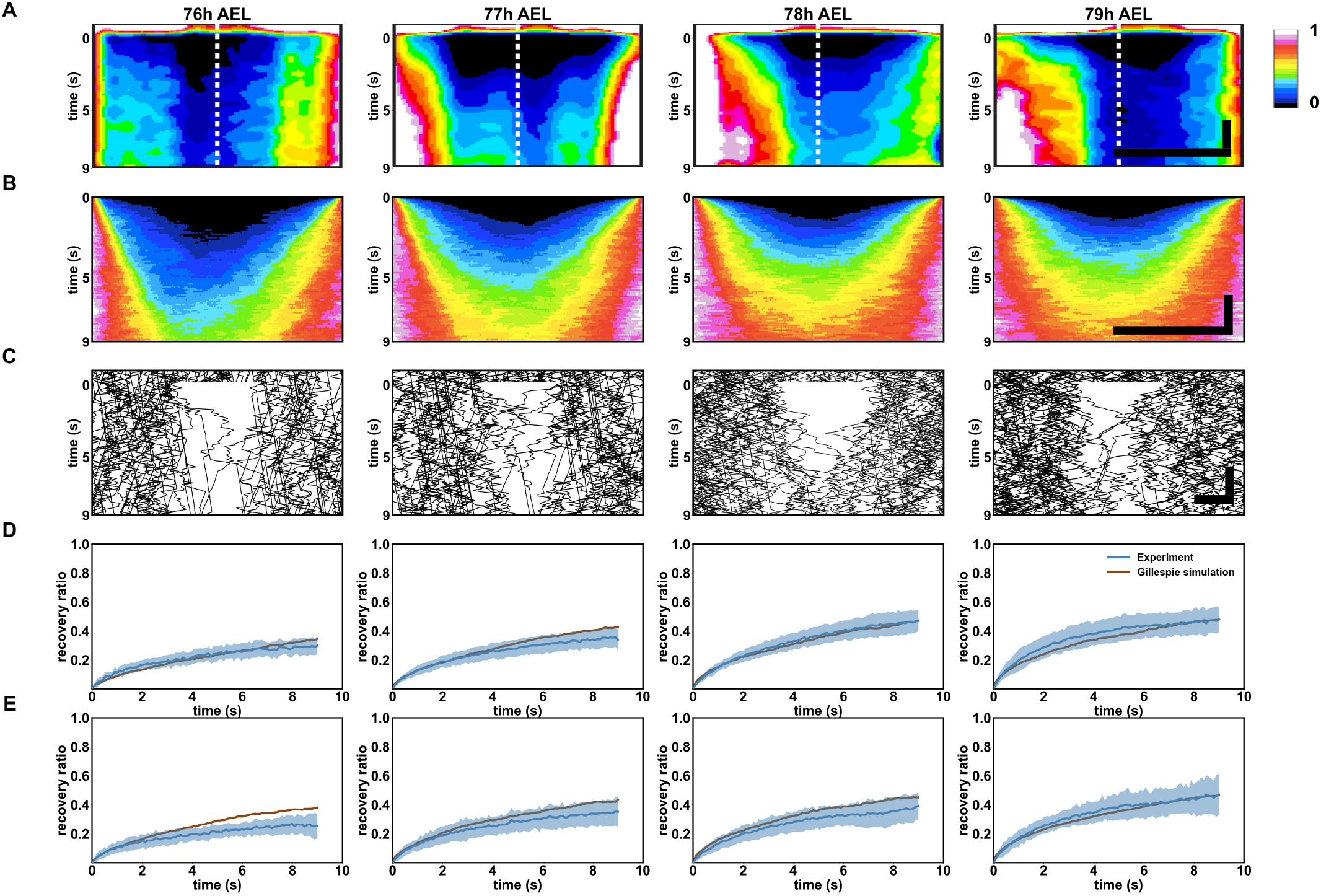
Results for the three state model with a diffusing state:(A-B)Colour kymographs of total cargo densities in the bleached region over 9 seconds from (A) experiment [(20) Image reproduced with permission from ..] and (B) Gillespie simulations for stages 76 - 79h AEL, (C) line kymographs over the entire region simulated using Gillespie algorithm for one run of the simulation, (D) proximal and (E) distal bleach recovery profiles from experiments and Gillespie simulations compared over 9 seconds. SCale bar - x-axis: 5*μm*, y-axis: 2*s*

The bleach recovery curves in case of the three state model with the diffusive state (Figure 4(D)) lie within the error bars of experimental recovery curves. They show a trend of rise in bleach recovery that is similar to the one obtained from experiments across these developmental stages. The simulation bleach recovery curves also show gradual rise followed by the flattening out that is seen in the experimental bleach recovery curves. Individual trajectories from one run of the simulations in Figure 4 C show the underlying transport behaviour producing the density colour kymographs in Figure 4 B.

We can extend this model by allowing for both a stalled state and a diffusive state. This increases the number of parameters that must be fit, including on and off rates that are currently unknown, such as the transition rate from stalled to either moving or diffusive. We have explored the consequences of such a model for completeness (Supplementary Information) but find that this, in fact, reduces the agreement slightly when compared to the simpler 3-state model with diffusion.

### Photoactivation

The influx of photoactivated GPAC-ChAT was measured as an increment in the GFP fluorescence in axon with respect to its preactivated form. Over the developmental time window of 76-78h AEL, the increment due to photoactivation revealed an episodic increase in GPAC-ChAT, mimicking the accumulation of GFP-ChAT in the Axonal Initial Segment (AIS) as well as the relative FRAP in the proximal halves. (Data for GFP-ChAT and FRAP are published in (20)). The photo-activated GFP-ChAT fluorescence shows the highest anterograde flow at 78h AEL. GPAC-ChAT accumulation in the AIS, measured by mCherry fluorescence, also showed an increasing trend from 76 to 78h AEL (Figure 5 B) corroborating the earlier results. This episodic increase during 76-78h AEL was not observed for soluble GFP which moves purely by diffusion and shows constant levels during these stages (Figure 5 A) in the AIS. Overall these results indicate that an episodic flow and accumulation of ChAT results from different modalities of soluble transport facilitated by Kinesin-2.

**Figure 5:**
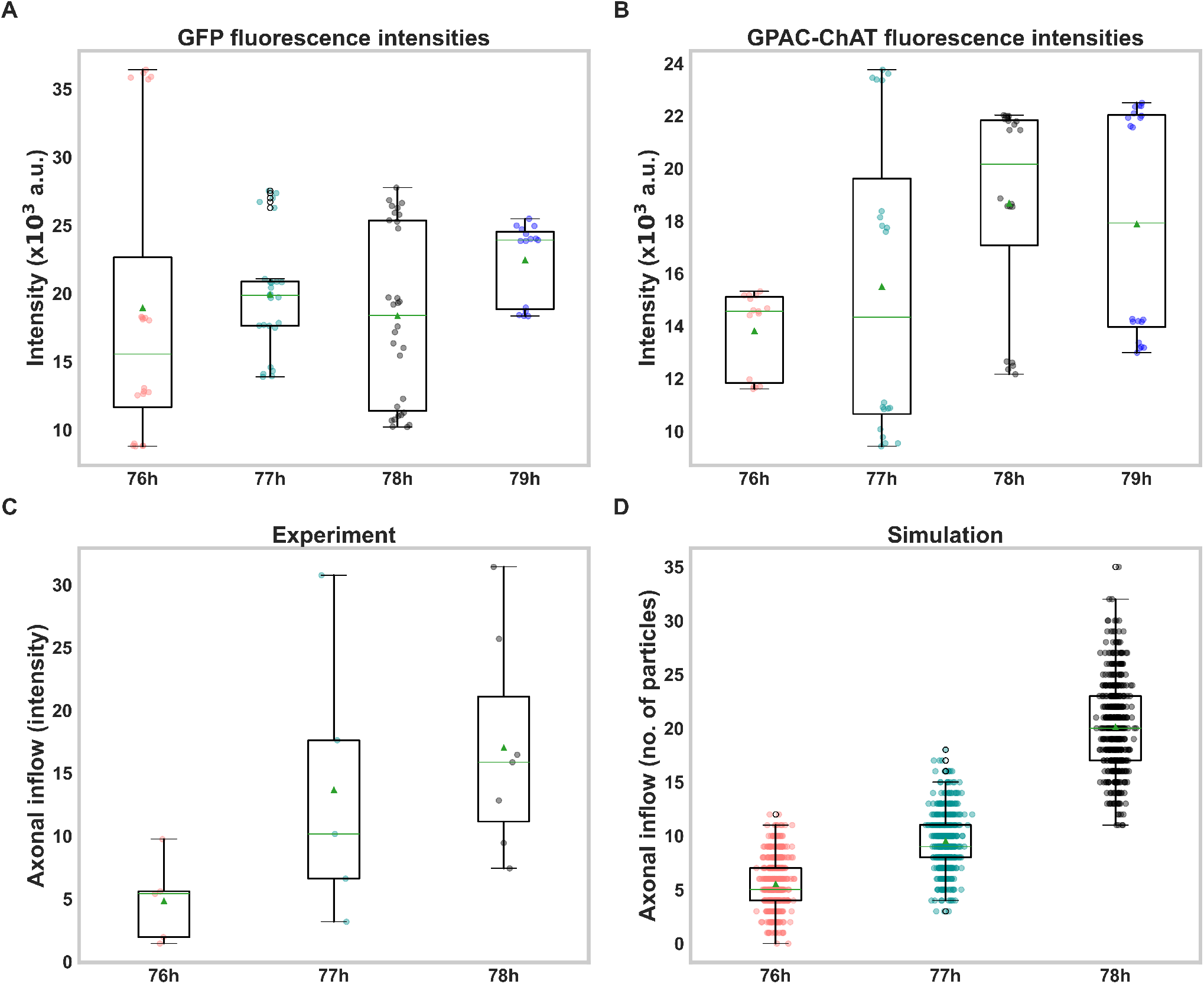
Fluorescence intensities of (A) GFP and (B) GPAC-ChAT in the axonal initial segment for stages 76 - 79h AEL. (C) Axonal inflow of GPAC-ChAT measured from 76-78h AEL as intensity increment due to photoactivation normalized to respective pre-activation intensities. (D) Cargo inflow from the cell body into 5*μm* region of the AIS following Photoactivation from Gillespie simulations (expressed as number of particles) for stages 76 - 78h AEL

To test the model, the photo-activation experiment were simulated with parameter values benchmarked as for photobleaching simulations. The amount of photo-activated cargo recovered in the 5*μm* region of interest 1s after photo-activation of cargo in the proximal region is shown in Figure 5 (D). Figure 5 (C) shows an increase in the amount of photo-activated cargo in the 5*μm* proximal AIS from 76 - 78h AEL observed in the experiments. A similar trend is seen in the simulation (Figure 5 D).

The amount of photo-activated cargo accumulating in the region of interest depends on the total cargo in the system. Two sources of measurements are available for the amount of cargo from experiments - TQ-ChAT and GFP-ChAT intensities (Figure S3 A). While the TQ-ChAT fluorescent intensities increases from 78 to 79h, GFP-ChAT intensities decreases in the same duration (Figure S3 A). Thus, using TQ-ChAT intensities to indicate the total cargo level in the system produces an axonal inflow trends that increases from 78 to 79h (Figure S3 C) while that with GFP-ChAT shows a minor decrease from 78 to 79h AEL (Figure S3 D). In the comparison with the photo-activation data we thus omit the 79h AEL (Figure 5 D).

## CONCLUSION

We have presented a model for the axonal transport of a molecule known to be transported via slow axonal transport mechanisms. Our model addresses experimental data on bleach-recovery measurements of the transport of the small, soluble protein Choline Acetyltransferase (ChAT) in lateral chordotonal (lch5) neurons of *Drosophila*. The model incorporates switching between paused, diffusive and directed transport. It is carefully benchmarked using available experimental data and biophysical estimates and studied using kinetic Monte Carlo simulations.

In the experiments, across developmental stages 76-78h AEL, there is a systematic increase in the amounts of cargo, motors and in cargo-motor interactions, as shown in Fig. 1. All of these must be factored into the calculation of how bleach recovery profiles vary across developmental stages. These are incorporated into the parameters in our model. Doing so leads to our main result – the increase in cargo overcompensates the increase in both motor numbers and in motor-cargo interactions, allowing a larger fraction of cargo to be transported by passive diffusion. Accounting for such diffusion leads to an overall increase in bleach recovery across these developmental stages, yielding bleach recovery curves in good agreement with the data. As a check on the modeling, we performed photoactivation measurements, showing that we could recover those results independently.

Several recent reviews (32) summarize insights from experiments (11) and modelling (33) related to the problems we study here. Smith and Simmons (25) first developed a “reaction-diffusion-transport” model with three cargo states for transport in axons governed by partial differential equations for diffusing and motor-driven cargo particles. We use the steady-state forms of these coupled equations to arrive at the relation between system parameters. Similar studies have been carried out by Brown et al (14) to study the slow transport of neurofilaments to propose the “stop-and-go” hypothesis for the transport. The model was further studied analytically by Jung and Brown (34). Experimental work on a class of SCb proteins has indicated transient assembly of proteins on fast moving cargo, leading to the proposal of the “dynamic recruitment” model for SCb transport. Studies of the transport of inactive Dynein showed a possible motor-limited scenario of transient association with Kinesin, with the intervening diffusion or pausing of the SCb component producing overall slower rates of transport. Theses studies point to the behaviour of the SCb proteins in the intervening period between the active excursions to play an important role in determining the rates of transport of these proteins. Other theoretical studies have modified the reaction-diffusion-transport model of Smith and Simmons to model the transport behaviour of specific SCb components. Kuznetsov et al (35) extend the model proposed by Brown and collaborators for the slow axonal transport of soluble proteins by including a diffusive state. Kuznetsov (36) also studies the transport of tau proteins in photobleach experiments using a model with two populations of tau - microtubule bound paused state and a diffusing free state and gaining insights on the behaviour of the two populations at the boundaries. Kuznetsov *et al* (37, 38) have studied mechanisms of slow transport of tau in axons. These studies solve the coupled differential equations for cargo in three and five states to describe the system numerically. However, predicting kymographs from such an averaged description is impossible.

Lattice models have been used to study the traffic of molecular motors (39–41) and cargo (42, 43) in various geometries - unbounded geometries (44), in open and closed compartments (45), through open tube-like compartments (46), half open tubes (47). The random walk exhibited by the molecular motors are modelled alternating between directed movement on filament tracks and diffusion in the surrounding cytoplasm. Ebbinghaus and Santen (40) study the transport of two particle species on two parallel lanes for directed movement with exclusion interactions and unbiased diffusive movement respectively. We do not model exclusion interactions in our simulation as interactions are not expected to play a major role in the transport of SCb proteins owing to their weak association with the motor and smaller sizes. Where our work differs from earlier model approaches is in our attempt to see what models and parameters could best describe data from experiments on a specific protein known to be transported via an SCb route, and in the detailed benchmarking of theory to experiment.

The methodology we outline and benchmark against data here improves on our understanding of the determinants of slow axonal transport in one, well-studied experimental system. A significant contribution of our work is that we are able to assess changes in the relative importance of different transport states across developmental stages, for which we know of no prior models. We also show that models that assume a stalled or paused state alone in addition to the transport states cannot reproduce the data on bleach recovery. Since we follow the trajectories of each cargo independently in our simulations, we can generate kymographs of individual protein trajectories as well as a number of other quantities that can be compared directly to experimental data. We show that incorporating a diffusive state reproduces properties of kymographs of particle trajectories across developmental stages 76 - 79h AEL. While we cannot rule out the possibility of a paused state (the more general 5-state model we study includes both paused and diffusive states), its relevance appears to be minor in these developmental contexts, since the data is very largely explained through the incorporation of the directed transport and diffusive states alone.

We expect that the study reported here should be relevant to our broad understanding of systems exhibiting slow axonal transport.

## AUTHOR CONTRIBUTIONS

Conceptualization: KR and GIM; Methodology: RM and GIM; Formal Analysis and Validation: RM, KR and GIM; Investigation: RM and SD; Writing: RM, GIM; Original Draft: all authors; Writing - Review and Editing: all authors; Visualization: RM and SD; Software: RM; Supervision: GIM; Project Administration: GIM

## SUPPORTING CITATIONS

References (20) appears in the Supporting Material.

## SUPPLEMENTARY MATERIAL

Document S1. Supplementary Information.

