## Supplementary Information for "The interplay of active and passive mechanisms in slow axonal transport"

### I Five state model

The five state model accounts for the following cargo motion states: (I) an anterograde motor-bound active state, (II) a retrograde motor-bound active state and (III) diffusive states, of which there are three. The diffusive states are: (i) a diffusive state with no motor bound, (ii) a paused state with anterograde motor bound and (iii) a paused state with a retrograde motor bound. The diffusing but motor-bound states can surrender their motor or motors to become purely diffusive. Also, motor-bound diffusive states can bind to microtubules and move in the anterograde or retrograde direction, depending on which motor is bound to the cargo.

As in the case of three state model (Figure 2) in the five state model, in an infinitesimal time interval, a particle can undergo one of the following: A particle in the active anterograde state can move one site to the right, at a rate $\gamma_h^+$ (Figure 2 B(ii)), while a particle in the active retrograde state can move one site to the left, at a rate $\gamma_h^-$(Figure 2 B(iv)), where the subscript $h$ indicates a hopping rate. A particle in the diffusive states can move one site to the right or left with equal probability $\gamma_D$(Figure 2 B(iii)). This rate is adjusted so as to yield a macroscopic diffusion constant that is characteristic of a diffusive particle of the same size as the protein. Additionally in the five state model, a particle not bound to a motor can associate with anterograde or retrograde motor at rates $\gamma_a^+, \gamma_a^-$ respectively and remain paused (Figure S1). In the paused state, the diffusion constant is set to zero (states DA and DR in Figure S1). A motor-bound paused particle can lose the associated motor to become a free diffusing particle at rates $\gamma_d^+, \gamma_d^-$ for anterograde and retrograde motor-bound diffusing particles respectively (Figure S1). A motor-bound paused particle can also bind to a microtubule at a rate $\gamma_b^\pm$, thus becoming an anterograde or retrograde mover depending on the nature of the bound motor (Figure S1). Likewise, an anterograde or retrograde particle moving on a microtubule can become a motor-bound paused particle by leaving the filament at a given rate $\gamma_u^\pm$ (Figure S1). The fundamental rates used in our simulation are listed in Table S1.

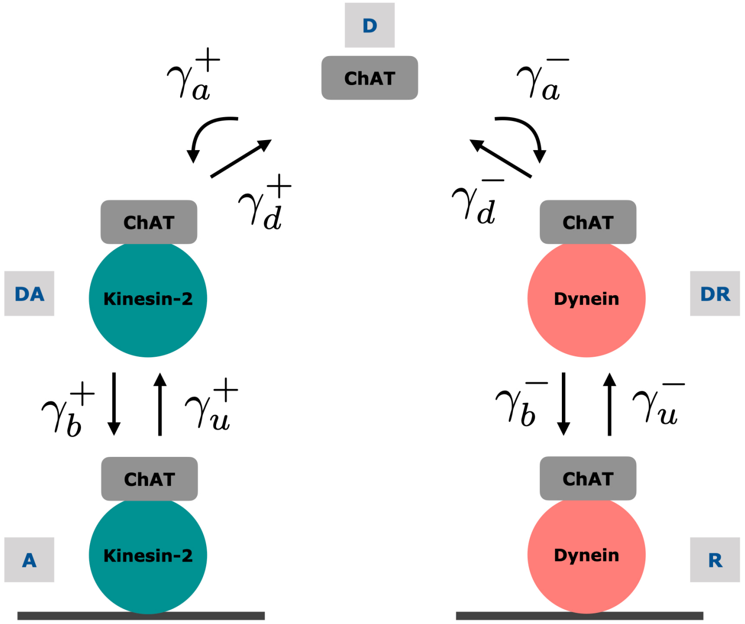


FIG. S1: **Schematic of the five state model for motor-driven axonal transport, depicting 5 states for the motor-cargo complex and transitions between states with the associated rates, These are described in the text.**

TABLE S1: **Parameters specific to 5 state model**

| **Symbol** | **Parameter** | **76h AEL** | **77h AEL** | **78h AEL** | **79h AEL** |
|---|---|---|---|---|---|
| $\gamma_b^+(s^{-1})$ | rate of kinesin-2-ChAT complex binding to microtubule track | 11.7 | 1.5 | 0.49 | 0.29 |
| $\gamma_b^-(s^{-1})$ | rate of dynein-ChAT complex binding to microtubule track | 30.22 | 4.34 | 1.26 | 0.86 |
| $\gamma_a^+(s^{-1})$ | rate of association of kinesin-2 and ChAT | 0.08 | 0.13 | 0.15 | 0.13 |
| $\gamma_a^-(s^{-1})$ | rate of association of dynein and ChAT | 0.08 | 0.1 | 0.1 | 0.1 |
| $\gamma_d^\pm(s^{-1})$ | rate of dissociation of kinesin-2 or dynein from ChAT | 1 | 1 | 1 | 1 |

For the five state model, the equations for steady-state are $\gamma_b^+\rho^{DA} = \gamma_u^+\rho^A, \gamma_b^-\rho^{DR} = \gamma_u^-\rho^R$, $\gamma_a^+\rho^D + \gamma_u^+\rho^A - (\gamma_u^+ + \gamma_b^+)\rho^{DA} = 0$, $\gamma_a^-\rho^D + \gamma_u^-\rho^R - (\gamma_b^- + \gamma_d^-)\rho^{DR} = 0$ and $\gamma_d^+\rho^{DA} + \gamma_d^-\rho^{DR} - (\gamma_a^+ + \gamma_a^-)\rho^D = 0$. Here, $\rho^A, \rho^R, \rho^D, \rho^{DA}, \rho^{DR}$ are densities of cargo in active anterograde, active retrograde, free diffusive, anterograde motor-bound paused and retrograde motor-bound paused states respectively.

Normalizing densities to the total cargo $\rho^T$ gives $\frac{\rho^{DA}}{\rho^T} = x$, $\frac{\rho^{DR}}{\rho^T} = y$, $\frac{\rho^D}{\rho^T} = r$, $\frac{\rho^A}{\rho^T} = p$ and $\frac{\rho^R}{\rho^T} = q$. We then have $x = (\frac{\gamma_a^+}{\gamma_d^+})r, y = (\frac{\gamma_a^-}{\gamma_d^-})r, p = (\frac{\gamma_b^+}{\gamma_u^+})x, q = (\frac{\gamma_b^-}{\gamma_u^-})y$. The unknown rates in the model are thus obtained: $\gamma_b^+ = \frac{\gamma_u^+ p}{x}, \gamma_b^- = \frac{\gamma_u^- q}{y}, (\frac{\gamma_a^+}{\gamma_d^+}) = \frac{x}{r}, (\frac{\gamma_a^-}{\gamma_d^-}) = \frac{y}{r}$.

**Results - Bleach recovery profiles across developmental stages**

Systematically varying the cargo distribution in the five cargo states around the distribution found to show better fits to bleach recovery profiles for the three state model, we arrive at a distribution that shows good fits to bleach recovery profiles across developmental stages 76 to 79h AEL (Table S2).

TABLE S2: **Cargo distribution (%) in 5 states for the parameter showing the best fit to experimental recovery curves**

| **Developmental Stage** | **76h AEL** | **77h AEL** | **78h AEL** | **79h AEL** |
|---|---|---|---|---|
| **Active anterograde (p)** | 45 | 29 | 17 | 10 |
| **Active retrograde (q)** | 40 | 23 | 10 | 8 |
| **Free diffusive (r)** | 13 | 39 | 58 | 66 |
| **Anterograde motor paused (x)** | 1 | 5 | 9 | 9 |
| **Retrograde motor paused (y)** | 1 | 4 | 6 | 7 |

Total fluorescent recovery in the proximal and distal bleached regions over time is plotted as the fraction of the pre-bleach fluoroscence in the region in Figure S2(D) across the developmental stages 76h - 79h AEL with error bars for the experimental profiels. The fits to proximal bleach recovery curves improve for 77 - 79h AEL with the inclusion of the paused states in the model, but the fits worsen for 76h AEL. The fraction of retrograde cargo in the paused and active states can in principle be adjusted to better fit the bleach recovery profiles. But since exact levels of retrograde motor and retrograde motor-cargo interactions are not know in the experimental system, we do not attempt to fit the distal recovery curves.

Bleach recovery profiles across the developmental stages 76h - 79h AEL from the Gillespie simulation (Figure S2(B)) are shown along with the experimentally derived colour kymographs (Figure S2(A)) from [1]. In Figure S2(C), we show particle kymographs across a larger region for one run of the simulation including the bleached region as a subset.

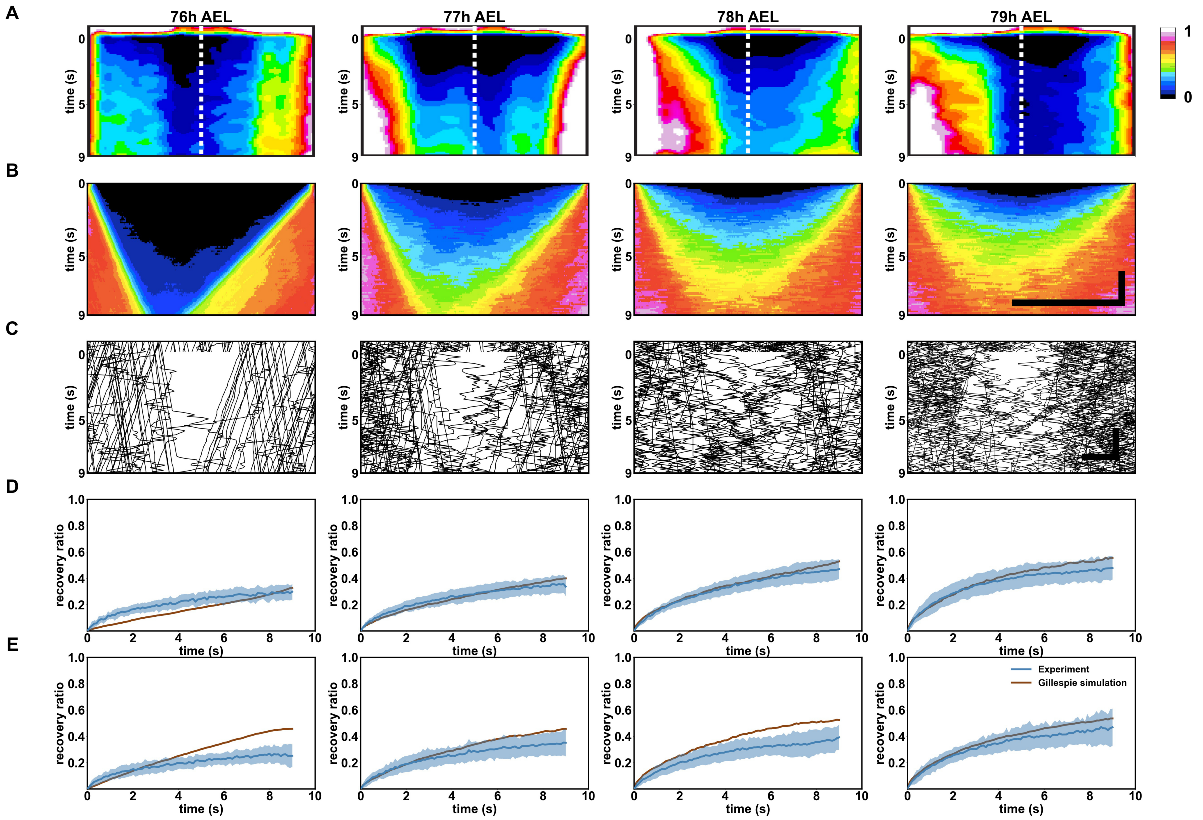


FIG. S2: **Results for the five state model with bulk diffusivity of the free diffusing cargo of $2.4\mu m^2/s$ and motor-bound cargo of $0\mu m^2/s$: (A-B)Colour kymographs of total cargo densities in the bleached region over 9 seconds from (A) experiment and (B) Gillespie simulations for stages 76 - 79h AEL, (C) line kymographs over the entire region simulated using Gillespie algorithm, (D) proximal and (E) distal bleach recovery profiles from experiments and Gillespie simulations compared over 9 seconds. Scale bar - x-axis: $5\mu m$, y-axis: $2s$**

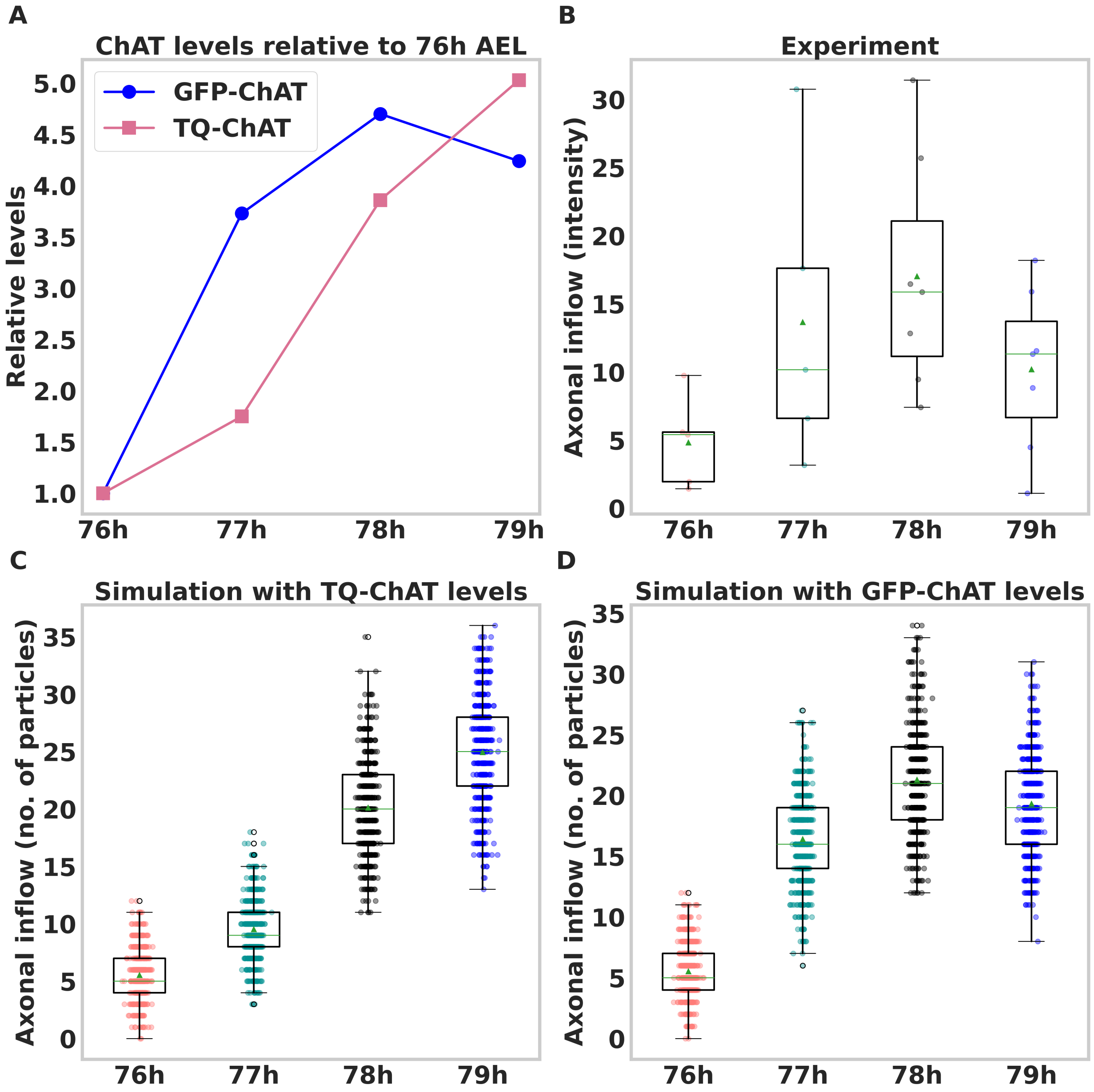


FIG. S3: **(A) Fluorescent intensity levels at developmental stages 76 to 79h AEL relative to levels at 76hAEL, (B) Axonal inflow of GPAC-ChAT measured from 76-79h AEL as intensity increment due to photoactivation normalized to respective pre-activation intensities. Cargo inflow from the cell body into 5$\mu m$ region of the AIS following Photoactivation from Gillespie simulations (expressed as number of particles) for stages 76 - 79h AEL using (C) TQ-ChAT and (D) GFP-ChAT fluorescent intensity measurements for total cargo in the system.**

[1] S. Dey and K. Ray, Scientific reports **8** (2018).

TABLE S3: **Cargo distribution (%) in 5 states for the parameter showing the best fit to experimental recovery curves**

| **Quantity** | **76h AEL** | **77h AEL** | **78h AEL** |
|---|---|---|---|
| TQ-ChAT levels relative to 76h AEL | 1 | 1.74 | 3.95 |
| Total cargo assumed | 50 | 87 | 197 |
| KLP68D levels | 1 | 1.14 | 1.48 |
| Fraction of total cargo taken to be active anterograde (changing across 76-79h AEL following the trend in KLP68D levels) | 0.35 | 0.23 | 0.13 |
| Fraction of cargo in active retrograde state (amount of cargo assumed constant across 76h-79h AEL) | 0.25 | 0.14 | 0.06 |
| Fraction of diffusive cargo (Total cargo minus amounts of active anterograde and retrograde cargo) | 0.4 | 0.63 | 0.81 |
